# Focused natural product elucidation by prioritizing high-throughput metabolomic studies with machine learning

**DOI:** 10.1101/535781

**Authors:** Nicholas J. Tobias, César Parra-Rojas, Yan-Ni Shi, Yi-Ming Shi, Svenja Simonyi, Aunchalee Thanwisai, Apichat Vitta, Narisara Chantratita, Esteban A. Hernandez-Vargas, Helge B. Bode

## Abstract

Bacteria of the genera *Photorhabdus* and *Xenorhabdus* produce a plethora of natural products to support their similar symbiotic lifecycles. For many of these compounds, the specific bioactivities are unknown. One common challenge in natural product research when trying to prioritize research efforts is the rediscovery of identical (or highly similar) compounds from different strains. Linking genome sequence to metabolite production can help in overcoming this problem. However, sequences are typically not available for entire collections of organisms. Here we perform a comprehensive metabolic screening using HPLC-MS data associated with a 114-strain collection (58 *Photorhabdus* and 56 *Xenorhabdus*) from across Thailand and explore the metabolic variation among the strains, matched with several abiotic factors. We utilize machine learning in order to rank the importance of individual metabolites in determining all given metadata. With this approach, we were able to prioritize metabolites in the context of natural product investigations, leading to the identification of previously unknown compounds. The top three highest-ranking features were associated with *Xenorhabdus* and attributed to the same chemical entity, cyclo(tetrahydroxybutyrate). This work addresses the need for prioritization in high-throughput metabolomic studies and demonstrates the viability of such an approach in future research.

---

*Photorhabdus* and *Xenorhabdus* are soil dwelling bacteria that are found worldwide in association with nematodes of the genera *Heterorhabditis* and *Steinernema*, respectively^1,2^. The bacteria live in symbiosis with their cognate nematode species and their life cycle involves a pathogenic stage towards invertebrate insects^3^. Although members of different genera, *Xenorhabdus* and *Photorhabdus* produce a number of shared specialized metabolites (SMs) and occupy very similar ecological niches^4^. Interestingly, the bacteria have yet to be isolated from the environment as free-living organisms, but instead are always found in association with their respective nematodes. Despite this specificity towards a nematode host, bacteria-nematode pairs may be isolated from the same geographic location.

Recently we highlighted the extensive chemical diversity present in these genera using high-throughput genomic and metabolomic analyses. It appears that SMs make up a major part of those coding sequences that were acquired and maintained in the genera upon divergence from a common ancestor, namely, members of the Enterobacteriaceae. We proposed that SMs, specifically products of polyketide synthases (PKSs) and non-ribosomal peptide synthetases (NRPSs), may be related to the given ecological niche that each strain occupies^4^. The products of these enzymes in *Photorhabdus* and *Xenorhabdus* have a range of known functions including antibiotic, signaling and assisting in development of the nematode host, among others (for recent reviews of all known natural products from these genera see ^5,6^).

One argument supporting an ecological function for the SMs, is the fact that although a few compounds appeared at first to be genus-specific, continued investigations have identified the same clusters in the other genus. Several clear examples of this are xenocoumacin, whose gene cluster was recently found in *Photorhabdus luminescens* PB45.5^7^ and xenorhabdin, whose gene cluster has been found in *Photorhabdus asymbiotica* strains^8^. Natural product research is continually encountering the problem of the best way to prioritize research efforts relating to “new” metabolites. One common way to do this is to find “new” genera or species that often produce a new subset of SMs^9^. Using genomic information to identify biosynthetic gene clusters that often produce bioactive compounds, such as PKSs or NRPSs, and subsequently activating “silent” clusters to specifically stimulate production of the metabolite in another way. However, in the absence of genetic information, this becomes increasingly difficult. Tools such as GNPS^10^, Sirius ^11^, MZmine^12,13^, DEREPLICATOR+^14^ and others have recently been developed for dereplication of MS/MS data. These have also been linked to several databases, which can assist in quickly identifying compounds absent in these databases. However, prioritizing the continued research and development of these unexplored metabolites is still a major problem.

Here, we describe the use of a machine learning model in order to explore the metabolomes of geographically distinct strains of *Photorhabdus* and *Xenorhabdus* from different regions in Thailand. We explored metabolic potential in relation to the environment in which they were collected, identified known compounds and prioritized the structure elucidation of one of the metabolites whose presence was most determining in distinguishing *Xenorhabdus* from *Photorhabdus*. Despite a number of long-standing hypotheses suggesting that metabolite production is specific to each strain (and its respective environment), this is the first time it has been empirically tested.

## Results

### Strain collection and processing

Strains selected for this study were collected from a variety of areas across central Thailand (Figure 1, Supplementary Table S1). Following isolation of the bacteria, each species was identified by sequencing and alignment of the *recA* coding sequence to the NCBI database (see Supplementary Table S1 for NCBI accession numbers). Our aim was to explore as big a metabolite repertoire as possible. We therefore cultivated in two different media; LB (nutrient rich) and SF900 (an insect-like medium), extracted each culture independently and combined the final results. Methanol was used to extract the cultures directly in equal volumes, which provided a robust dataset on which to perform further analyses. Acetonitrile blanks and media only were used to subtract background masses while *E. coli* (a close relative of *Xenorhabdus* and *Photorhabdus*) was additionally used in order to determine metabolites that were not likely specific to the *Xenorhabdus* and *Photorhabdus.* The combined analysis identified a total of 44,836 molecular features after removing background features (LB, SF900, acetonitrile and *E. coli* in both media). MS data sets can be found under public MassIVE ID: MSV000083378 and the combined network analysis can be downloaded at http://gnps.ucsd.edu/ProteoSAFe/status.jsp?task=02057a6b9eb54048847c9dd18746aac9.

**Figure 1a.**
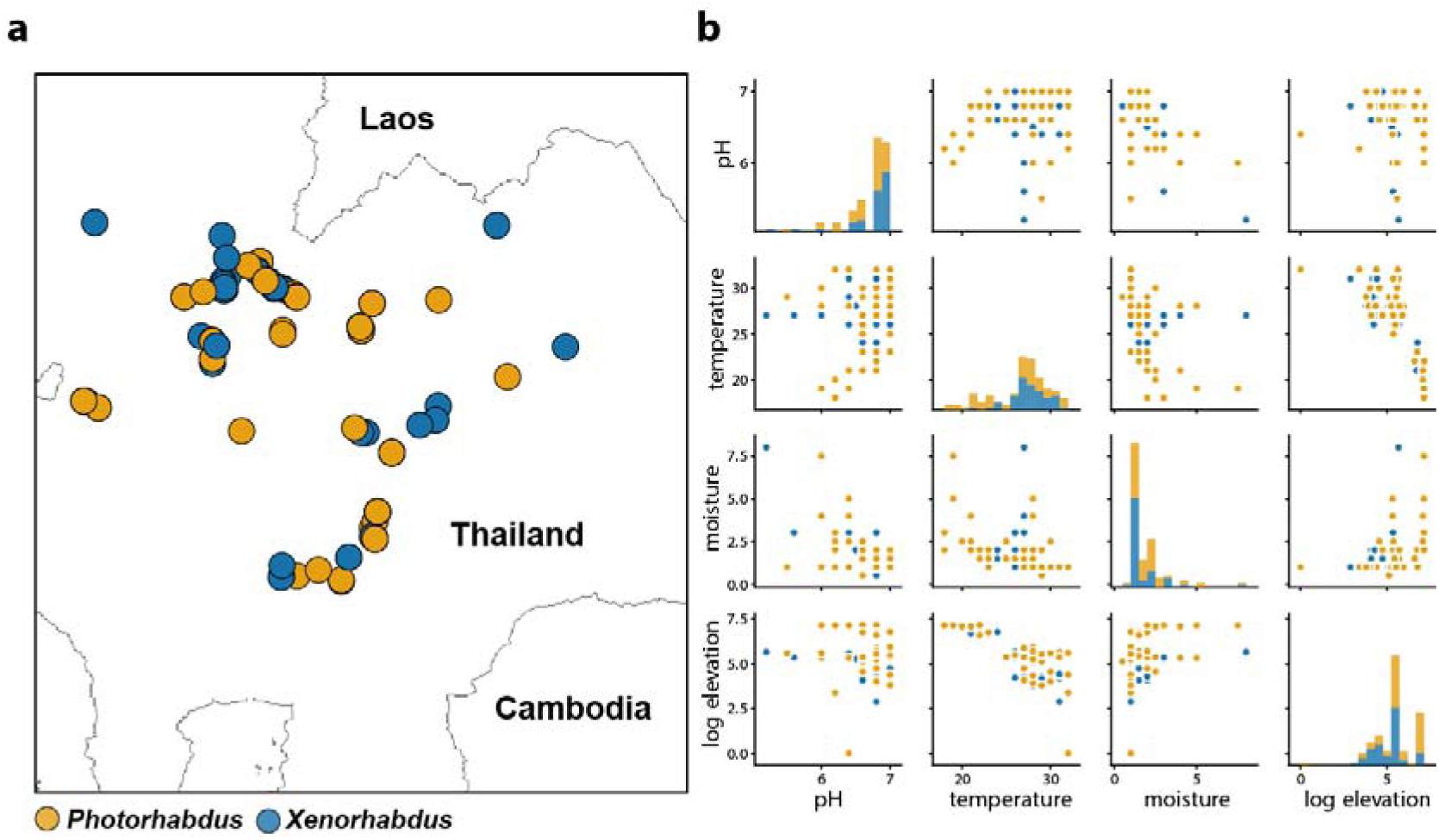
Location and b) spread of metadata associated with the 114 *Photorhabdus* and *Xenorhabdus* strains collected from Thailand. For specific metadata values, see Supplementary Table S1.

### Network analysis

Network analysis was performed on the complete collection of strains using GNPS^15^ and Cytoscape^16^ for visualization (Figure 2). *Xenorhabdus* had a greater number of unique molecular features (3,265), compared to the *Photorhabdus* (1,791). A total of 261 networks with three or more nodes were formed (Figure 2). Of these, 14 families of compounds could be identified based on previously published studies, leaving a majority of networks still completely unexplored. Use of GNPS and its resulting network analyses, revealed a number of networks containing known compounds. These networks group metabolites with structural similarity based on their fragmentation patterns^15^. We assume that all nodes within a given network belong to the same metabolite family. We have shown in *Photorhabdus* and *Xenorhabdus* that this is indeed often the case, as described in our previous work^4^. Despite providing a broader perspective on the presence and absence of metabolite families, what this fails to address is whether or not these nodes and/or metabolites are important in defining any variables that may be interesting for further investigation.

**Figure 2.**
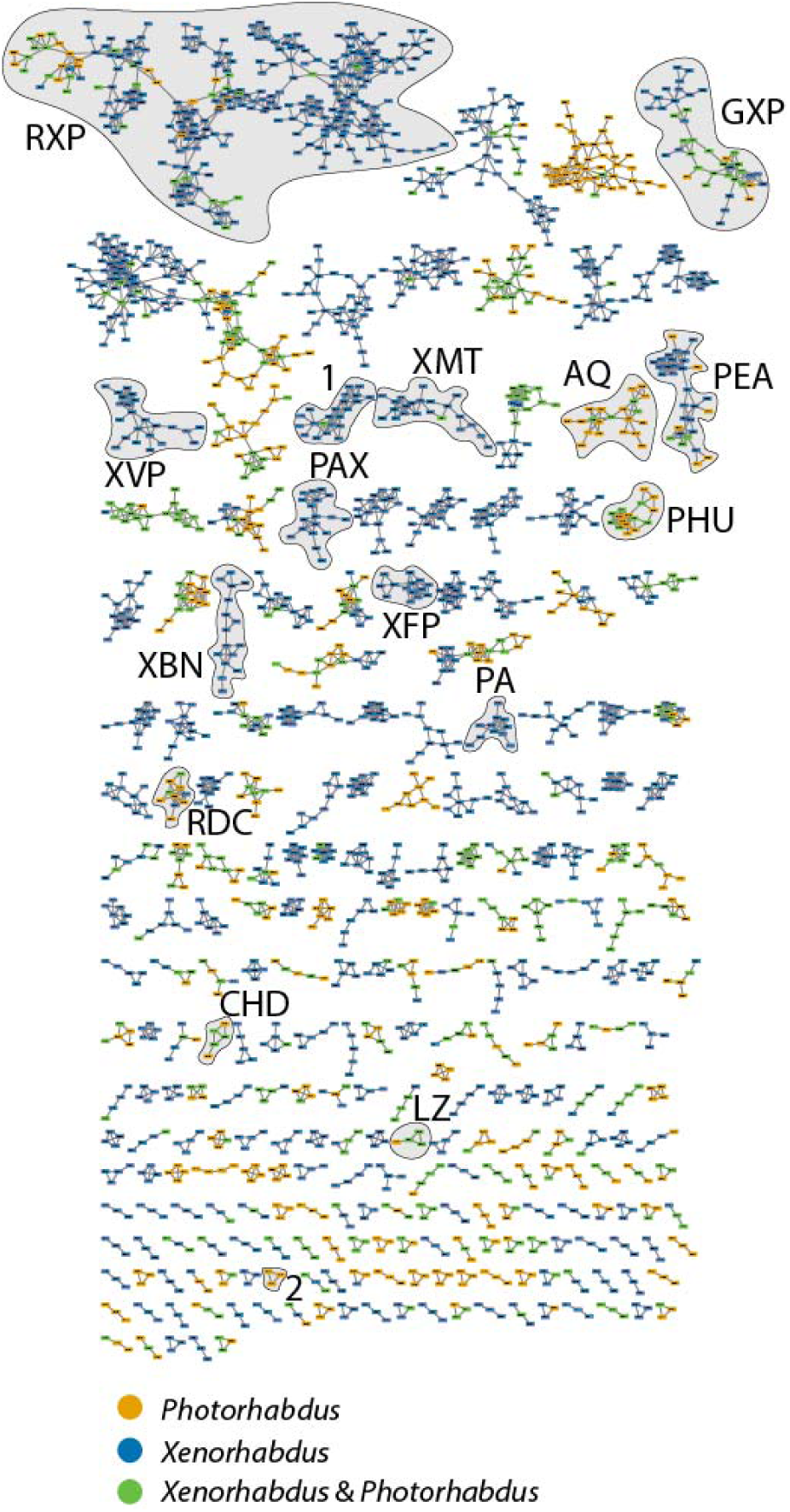
Network analysis of all 114 isolates. Shown is a summary of all nodes with at least two connections in *Photorhabdus* and *Xenorhabdus*. Known subnetworks are also highlighted: RXP – rhabdopeptide, GXP – GameXPeptide, XVP – xentrivalpeptide, PAX – PAX peptide, AQ – anthraquinone, PEA – phenylethylamide, XFP – xefoampeptide, CHD – cyclohexanedione, LZ – luminizone, RDC - rhabduscin, PA – pyrrolizidine alkaloids, XBN – xenobactins, XMT – xenematide/xenoprotide, **1** – (cyclo)tetrahydroxybutyrate, **2** – network containing signal with *m/z* of 487.18. For a closer view of the network containing **1**, see Supplementary Figure S3.

We have discussed at length the possibility for analogous functions by different *Photorhabdus-* or *Xenorhabdus-*specific compounds^4^, which would help explain the reasons they live such a similar lifestyle. However, as is clear from Figure 2, there is still a significant number of metabolite clusters yet to be explored. This begs the question as to where we should focus our research efforts in looking for unknown and important compounds, with respect to both the bacterial ecology and natural product discovery. We therefore decided to utilize machine learning in order to prioritize compounds and their investigations, with an end-goal of researching metabolites that are likely to be both undiscovered and specific.

### Machine learning to explain metadata

Our data consisted of a total of 114 different strains, coupled to seven abiotic metadata points; two media conditions, four soil types, ten provinces representing rough geographic relatedness, soil pH, soil temperature, soil moisture and elevation above sea level. In order to explore our data in more detail and determine what, if any, of these abiotic factors could be distinguished by utilizing metabolite production, we turned to machine learning. We utilized a gradient boosting decision tree algorithm in order to train the model on the full dataset, as well as a reduced dataset consisting of highly-correlated signals (see Methods).

Training the model on the full versus the pruned and clustered datasets (Supplementary Figure S1) results in essentially the same performance (Supplementary Table S2). An initial analysis failed to show any significant impact of the abiotic data on metabolite production. Additionally, both randomizing and removing geographical metadata from the dataset did not result in a performance drop. We incorporated SHapley Additive exPlanations (SHAP) values into our model in order to determine the importance of individual features on model output. For both AUC (area under the curve) and intensity datasets with low levels of clustering, we see that a small number of metabolites strongly affect the output of all samples, and seem to do so in a well-delimited fashion (Figure 3). The impact of a few others is not as strong, but retain the latter property.

**Figure 3.**
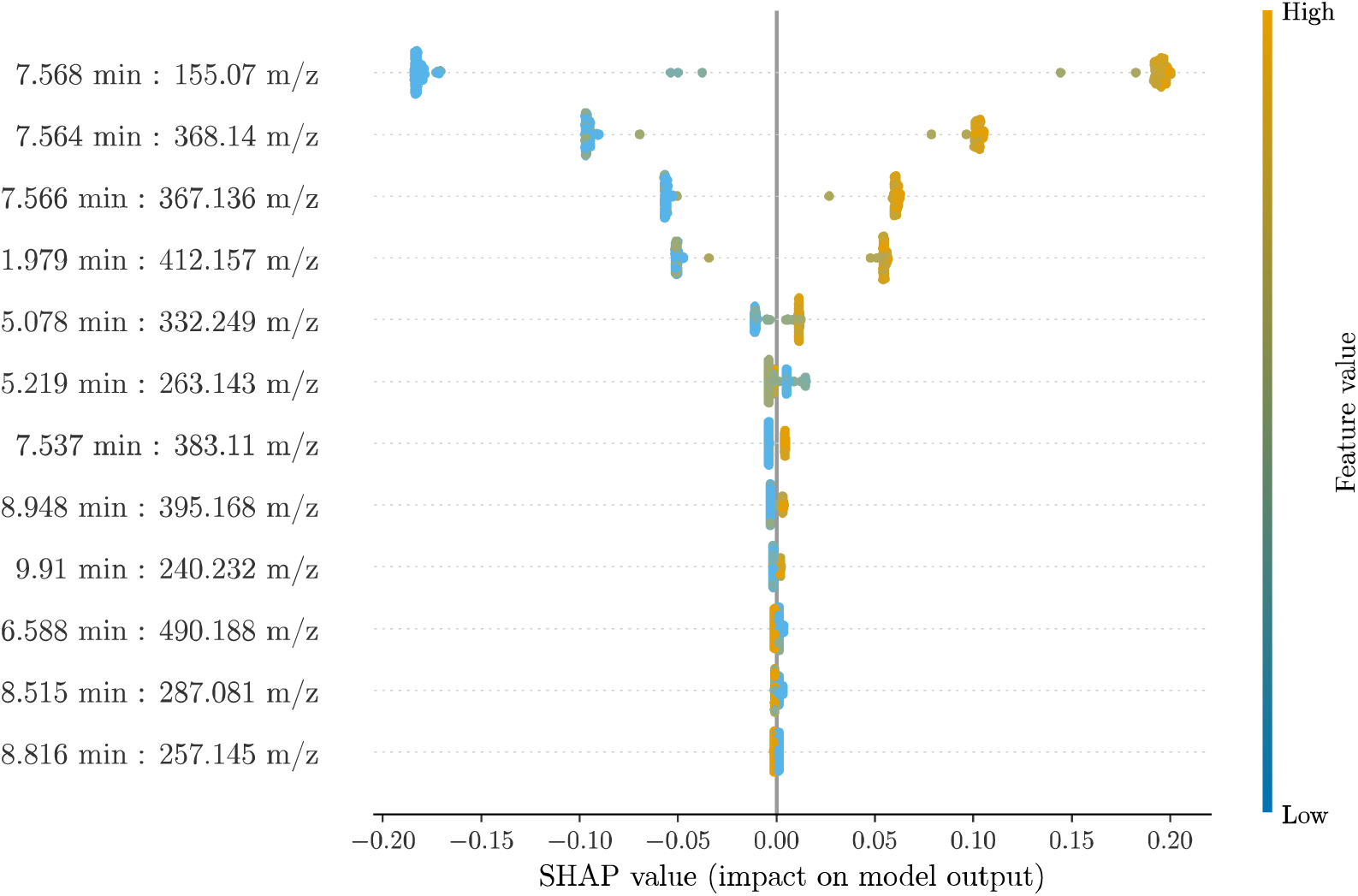
SHAP output of the GBDT model constructed using intensity values. The value represents the impact of a given feature in determining whether an isolate is *Photorhabdus* or *Xenorhabdus*. The *m/z* ratios and retention times are indicated for the top 10 ranking features.

### Structure elucidation of top-ranking feature(s)

Multiple metabolites seemed to be independently capable of discerning between genera with a high degree of accuracy. In particular, the top three single-feature predictors possessed the same retention times with *m/z* of 155.07, 368.14 and 367.13, respectively (Supplementary Figure S2). All three of these metabolites were highly correlated, with the third compound additionally identified in the network analysis (Figure 2, Supplementary Figure S3) and produced in large amounts in a strain of *X. szentirmaii* (see Methods).

Compound **1**, obtained as a colorless crystal, has the molecular formula C_16_H_24_NaO_8_ as deduced from its HR-ESI-MS at *m*/*z* 367.1366 [M+Na]^+^ (calcd for C_16_H_24_NaO_8_, 367.1363) in combination with ^1^H and ^13^C NMR data (Supplementary Table S3, Supplementary Figures S14-S18). By comparing its spectroscopic and single-crystal X-ray diffraction data with those reported previously in literature, it was identified as (4*R*,8*R*,12*R*,16*R*)-4,8,12,16-tetramethyl-1,5,9,13-tetraoxacyclohexadecane-2,6,10,14-tetrone, a cyclic tetramer of (*R*)-3-hydroxybutyrate (Figure 4, Supplementary Table S3)^17,18^. The presence of the signal with an *m/z* of 155.07 can also be explained by the structure of **1** (Figure 4c), while the signal with *m/z* of 368.14 is the ^13^C isotope of **1**.

**Figure 4.**
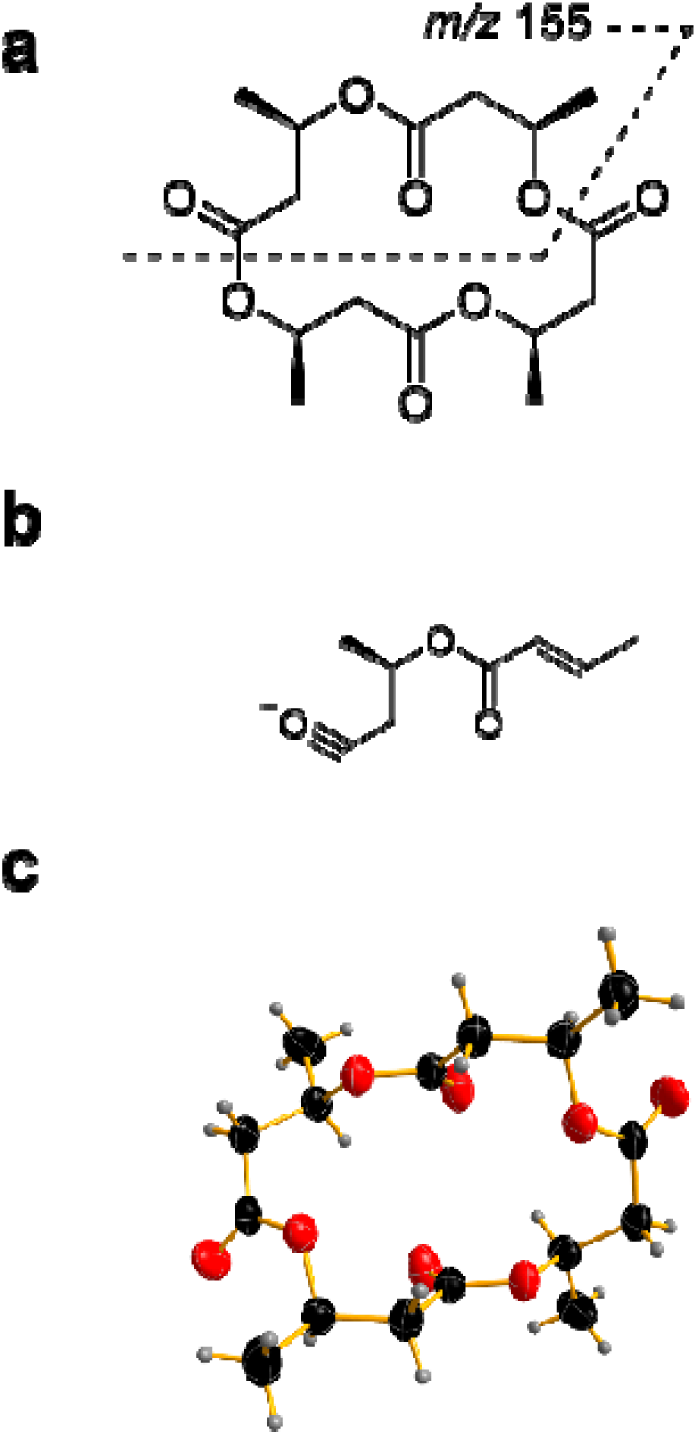
Structure of (4*R*,8*R*,12*R*,16*R*)-4,8,12,16-tetramethyl-1,5,9,13-tetraoxacyclo hexadecane-2,6,10,14-tetrone (1). The structure **(a)** and the fragment responsible for the signal at *m/z* 155 is indicated **(b)** as well as the ORTEP representation of its crystal structure (CCDC 1880748) **(c)**.

### Single features are capable of discerning genera with high accuracy

Higher clustering (lower correlation thresholds) of the metabolite data resulted in the signal with an *m/z* of 155.07, being identified as having, by far, the largest influence in model output in all cases (Supplementary Figure S4-S9). Focusing on all metabolites belonging to the same cluster as this metabolite, as well as those belonging to the clusters represented by the metabolites ranked second and third by SHAP values, we proceeded to retrain the model employing as a feature only one metabolite at a time. We found that the three best single predictors in terms of ROC-AUC (receiver operating characteristic – area under the curve) for both the intensity and AUC data corresponded to signals with an *m/z* of 155.07, 368.14, and 367.14 (Figure 3). These can be used as sole predictors while maintaining a very high performance, equivalent to using the full set of metabolites (Supplementary Table S4).

To explore whether the three top ranking features, all belonging to the same cluster of signals, significantly impacted the model’s performance, we removed all features associated with this cluster and recalculated the model. The resulting top-ranking feature and it’s highly-correlated features were again removed and the model recalculated a third time for comparison. The performance after removing these clusters remained high at 95.2% ± 1.44% and 95% ± 1.3%, respectively, with other signals showing a highly discriminatory effect between *Photorhabdus* and *Xenorhabdus* (Supplementary Figure S10 and S11). However, the top three clusters all related to features present in the *Xenorhabdus* and absent in *Photorhabdus.* We therefore identified features that were negatively correlated to the top-ranking cluster and used this as a sole predictor for the genera. In essence, the original model was able to predict a *Photorhabdus* by the absence of the three aforementioned top-ranking features. By using a negative correlation, we aimed to identify compounds that were present in a majority of *Photorhabdus*, but absent in *Xenorhabdus*. This resulted in the identification of a signal with an *m/z* of 487.19 (predicted sum formula: C_26_H_25_N_5_O_5_), whose fragmentation pattern suggests it might be a peptide (Supplementary Figure S12). Additionally, this metabolite was also detected in the network analysis, albeit in a much smaller cluster of nodes (Figure 2). Using this feature as a sole predictor of genus resulted in a performance of 92.9% ± 2.99%.

### Model testing on unseen data

Fourteen *Photorhabdus* and 15 *Xenorhabdus* were randomly selected from the strain collection used for generating the original model, grown and extracted from both media types, in triplicate. These new HPLC-MS runs, unseen by the model during training, were used to test its general performance. From the metabolites present in the data, we located the closest match (see Methods) for each of the three previously-identified best predictors and obtained the class probabilities for each sample. In all cases, the single-feature models were able to correctly classify the genera of the samples with 92.0%-96.5% accuracy. The results are summarized in Supplementary Table S5.

## Discussion

Typically, the similarities between *Photorhabdus* and *Xenorhabdus* are highlighted, particularly with respect to their life cycles. While these similarities hold true, several recent efforts have sought to decipher their differences and what makes these genera unique^4,19^. Our recent work approached this from more of a genomic perspective, while here we attempt to answer this same question using metabolomics as a guide.

It is known that *Photorhabdus* and *Xenorhabdus* are capable of infecting different insect species leading to profoundly different experimental outcomes. This is probably because of the number of compounds which, generally speaking, suppresses the innate insect immune response^5^. What we don’t know however, is the degree of dependence that the bacteria have upon their repertoire of metabolites to adapt to the abiotic environment. Interestingly, these bacteria have not yet been isolated as free-living organisms; only in conjunction with their cognate nematode symbionts. We wanted to explore the hypothesis that strains collected in geographically different and abiotically diverse environments (pH, soil type, soil temperature, soil moisture, elevation above sea level) produce different metabolites, specific to that environment, thereby maintaining some form of localized niche despite the mobility afforded by nematode hosts.

A large collection of *Xenorhabdus* and *Photorhabdus* strains was acquired from Thailand, including a number of samples collected from the same geographic locations (Figure 1). Once isolated, we hypothesized that, by growing the strains under different conditions and collating the data, we would have a data set that represented the metabolic potential of each of the 114 strains. For that reason, we grew the strains in a rich media (LB) in order to provide an environment whereby it would not be disadvantageous (from an energy perspective) to produce compounds and also in SF900, an insect culture medium that reflects the environment these strains may encounter within an insect. A network analysis of the 58 *Photorhabdus* and 56 *Xenorhabdus* was performed using the GNPS platform, which examines mass differences and fragmentation patterns between metabolites in order to determine whether they are likely to be related from a chemical perspective. Despite the over-representation of some species in this collection, a combined network analysis of the 114 strains in both media highlights the chemical diversity present in Thailand by entomopathogenic bacteria, regardless of species (6,890 nodes, Figure 2). Our previous work annotated a number of metabolites from both *Photorhabdus* and *Xenorhabdus* and using this library, we identified 14 networks containing known clusters of metabolites (Figure 2). It is also clear from these analyses that there are a number of major metabolite families that we have yet to identify. Furthermore, it is known that both *Photorhabdus* and *Xenorhabdus* have several different mechanisms at their disposal to help generate natural product diversity from a single gene cluster^20,21^. In fact, the rhabdopeptides are known to be virulence factors towards insects and have an unusual mechanism of generating SM variation by altering the stoichiometry of each module^20^. This variation may actually contribute to the ability of these bacteria to infect different insects, adapting to different insects primarily by altering protein expression levels. In this analysis, we see a large number of features (330) in the network containing known rhabdopeptides (Figure 2). If this is a major factor conferring virulence to the bacteria, this might be indicative of an insect-specific adaptation.

These bacteria are of general interest due to their SM producing abilities. A recent rarefaction analysis of all sequenced *Xenorhabdus* and *Photorhabdus* genomes suggests that sequencing of a new species would yield, on average, one additional biosynthetic gene cluster per species sequenced. Notably, a recent study in *Myxobacteria* highlights the fact that strain collections with a threshold of taxonomic diversity and coverage is required in order to rapidly identify compounds with a high likelihood of containing structural novelty^9^. In this analysis, there was a large over-representation of *X. stockiae* species, but several new derivatives of known compounds. While we don’t dispute that structural novelty is important, we do observe that natural structural diversity present in bacteria that make compound libraries may also be important for structure-function studies. To that effect, the generation of new derivatives of known SM from these bacteria, through *in vitro* combinatorial biosynthesis, is ongoing with a view to identifying compounds with higher bioactivities^22^. What our analysis suggests is that, there is a strong possibility that many of these derivatives may also exist “naturally” in the environment as evidenced by the extensive molecular networks containing “known” compounds. Despite the apparent abundance of new derivatives, this also suggests that our prediction of one new SM per species is a significant under-estimation if we consider unknown derivatives.

Recently it was found that genes in strains isolated from similar environments, which are also the same species, contain a number of differences at the genetic level^23^. We envisaged that we may therefore be able to differentiate between different metadata based upon each strain’s unique metabolome. We used the compiled metabolomic data, together with the metadata, to train a machine learning model; in particular, we chose to make use of gradient boosting decision trees (GBDTs). Models of this type enjoy a high level of popularity due to their high efficiency and state-of-the-art performance, as well as the availability of fast, ready-to-use implementations. In addition to this, they tend to perform well, even in very-high-dimensional scenarios, especially in cases when the features outnumber the samples or observations, a phenomenon commonly referred to as the “curse of dimensionality”^24,25^. As such, GBDT models are ideally suited for the type of data we are dealing with – and metabolomics data in general – having tens of thousands of metabolites for a few hundred samples.

In addition to the above, GBDT models are also robust to multicollinearity between features. As seen from the results, the model does not suffer a performance drop when highly-correlated metabolites are present. Nevertheless, we decided to cluster the metabolites, and drop correlated variables, for interpretability reasons: faced with two or more highly-correlated features that are very good predictors, the model will greedily choose to split on one of them in detriment of the others. In other words, features that are otherwise highly discriminatory will have their impact underestimated in the ranking of importance.

One weakness in studies such as this, is the use of artificial *in vitro* culture conditions to explore the metabolic diversity. In comparative genetic studies, we typically compare whole genomes to draw inferences on the data, thus basing future hypotheses on the genetic potential, rather than gene expression. In the same principle, we base our conclusions here on metabolic potential and work towards overcoming the limitations associated with the non-natural environment by using different conditions and collating the data. Given that no evidence was seen for metadata influencing metabolite production, we used a machine learning model to investigate the differences between *Photorhabdus* and *Xenorhabdus*. During training of the model, SHAP values were obtained in order to assess and rank the impact of the feature values on model output. Our reasoning behind this was that we could then prioritize metabolites for purification and chemical structure elucidation. We chose SHAP values as our measure of importance because they provide per-sample explanations which are proven to be both consistent and locally accurate, as opposed to GBDTs built-in measures^26,27^, in addition to being a model-agnostic feature attribution approach that does not require the model to be tree-based.

From the SHAP results we observe that, while only a few metabolites – exactly one, for the most heavily clustered data – has a very large impact on model output in comparison to the rest, many more seem to be strong discriminators between classes, as evidenced by the coloring of their values and the direction of their impact, despite the latter being relatively low. Indeed, removal of the most important cluster from the dataset still resulted in very high classification performance when taking all other metabolites in consideration (Supplementary Figure S10). Single-feature predictions, however, do suffer from a steeper performance drop compared to the metabolites we have identified as the best predictors. Therefore, we emphasize that we have not attempted to find the ‘only’ metabolites that set these two genera apart, but to prioritize the ones that appear to be the strongest in doing so. The relevance of this, and the usefulness of single-feature models, becomes apparent when dealing with new, unseen data: in the case presented here, the test dataset contains 15,098 metabolite columns, which renders futile any attempt at full dataset peak matching.

A recent study in Australia examined the differences between the biosynthetic domain compositions in soil across the continent. One key finding from this was that the composition of natural product domains, specifically ketosynthase domains (from PKS) or adenylation domains (from NRPS), changed with latitude and longitude and was often grouped in accordance with the vegetation type^28^. This supports our original premise that natural product composition from the *Xenorhabdus* and *Photorhabdus* may change within the country. However, in our analysis we saw no clear clustering of strains based on any of the abiotic factors measured. Considering that the bacteria have never been isolated independent of the nematode, several explanations exist for the lack of obvious metabolite clustering in different environments. One explanation is that the nematodes, and the insects that they infect, are all motile and may help spread the bacteria in the environment thus confounding any underlying association with geography. One further explanation is that the nematode hosts provide the greater support in these environments. In turn, the specialized metabolites produced by the bacteria then provide specificity for the host and the invertebrate prey. This would actually point towards a dependence of the bacteria upon the nematode in the environment, an area that has not been widely investigated due to the relative simplicity to investigate the bacteria independently in a lab environment.

Purification of compound **1** resulted in elucidation of a cyclic tetramer of hydroxybutyrate (Figure 4), a compound related to crown ethers. Crown ethers typically demonstrate a high affinity to cations and are often cytotoxic, but may also show characteristics of ionophores. Ionophores in natural biological systems help to transport ions across cell membranes by forming lipid-soluble complexes with polar cations^29^. Given the probable influence of nematode host on metabolite production, one explanation for the specific presence of these compounds in *Xenorhabdus* could be that they are required during the symbiosis with *Steinernema*. While this is probably not a ubiquitous requirement since the compound was not detected in all species of *Xenorhabdus* (Supplementary Figure S13), it is interesting that the majority of the *Xenorhabdus*, with the exception of *X. szentirmaii*, were originally isolated in South East Asia. One interesting note is that the nematode hosts of *X. szentirmaii* (*Steinernema rarum*) and *X. stockiae* (*Steinernema siamkayai*) are close evolutionary relatives^30^, supporting a possible role of this metabolite in symbiosis.

One major challenge in large-scale metabolomic studies is how to prioritize research efforts. Here, we set out an analysis pipeline that is capable of using strain-specific metadata, coupled to high-throughput MS experiments. Whether it is determining compounds important for an ecological niche or identifying as yet undiscovered compounds in large high-throughput screening experiments. By incorporating machine learning models such as this into current analysis pipelines, the relative importance of compounds can be determined in order to streamline purification and/or structure elucidation pipelines in a time-efficient manner, yielding low probabilities of rediscovery.

## Materials and Methods

### Soil collection

Samples were taken from diverse habitats including natural grassland, roadside verges, woodlands, and banks of ponds and rivers. For each site, 5 soil samples were randomly taken in an area of approximately 100 m^2^ at a depth of 10-20 cm using a hand shovel. Approximately 500 g of each soil sample was placed into a plastic bag. The longitude, latitude and altitude of each sampling site were recorded using a GPSMAP 60CSx (Garmin, Taiwan). The temperature, pH and moisture of each sample were recorded using a Soil pH & Moisture Tester (Model: DM-15, Takemura electric works, Ltd, Japan).

### Isolation of *Xenorhabdus* and *Photorhabdus* bacteria from entomopathogenic nematodes

Dead *Galleria mellonella* larvae were surface-sterilized by dipping into absolute ethanol for 1 min and placed in a sterile petri dish to dry. Sterile forceps were used to nip the 3^rd^ ring from the head of *G. mellonella*, thereby removing the cuticle. A sterile loop was used to touch haemolymph of *G. mellonella* and streaked onto a nutrient bromothymol blue agar (NBTA) supplemented with 0.004% (w/v) triphenyltetrazolium chloride (TTC, Sigma, St. Louis, KS, USA) and 0.0025% (w/v) bromothymol blue^31^. TTC was added to inhibit the growth of Gram-positive, acid-fast bacteria and actinomycetes. Cultured plates were incubated in the dark at room temperature for 4 days. *Xenorhabdus* and *Photorhabdus* strains were characterized based on colony morphology as described by Boemare and Akhurst^32^. Single colonies were then subcultured on the same medium and kept in Luria-Bertani (LB) containing 20% glycerol at −80°C for further identification.

### Bacterial identification

DNA was extracted using a Genomic DNA Mini Kit (blood/Cultured Cell) (Geneaid Biotech Ltd., Taiwan). Polymerase Chain Reaction (PCR) targeting *recA* was performed in 50 *µ*l volumes using 10 *µ*l of 5X buffer (Promega, Madison, WI, USA), 7 *µ*l of 25 mM MgCl_2_ (Promega, Madison, WI, USA), 1 *µ*l of 200 mM dNTPs (New England Biolabs Inc., Ipswich, MA, USA), 2 *µ*l of 5 *µ*M of each Primer, 0.5 *µ*l of 5 unit Taq Polymerase (Promega, Madison, WI, USA) and 2.5 *µ*l of DNA template. The *recA* primer sequences were recA1_F (5’-GCTATTGATGAAAATAAACA-3’) and recA2_R (5’-RATTTTRTCWCCRTTRTAGCT-3’)^33^.

PCR cycling parameters for *recA* of *Xenorhabdus* included an initial denaturing step of 94°C for 5 min, followed by 30 cycles of denaturation at 94°C for 1 min, annealing temperature of 50°C for 1 min and extension of 72°C for 2 min and a final extension of 72°C for 7 min. Parameters for *Photorhabdus* included an initial denaturing step at 94°C for 5 min, followed by 30 cycles of 94°C for 1 min, 50°C for 45 sec and 72°C for 1.5 min, with a final extension of 72°C for 7 min. The PCR products of *recA* of both genera (890 bp) were examined on 1.5% agarose gel electrophoresis. Fifty microlitres of PCR products were purified using Gel/PCR DNA Fragments Extraction Kit (Geneaid Biotech Ltd., Taiwan). *recA* sequencing was performed on the ABI PRISM® 3100 Genetic Analyzer (Amersham Bioscience, UK) using the PCR primers for PCR. Chromatograms, sequence ambiguity resolution were visually checked using SeqManII software (DNASTAR Inc., Wisconsin, USA). Species identification was performed using a nucleotide Blast search of *recA* against the NCBI nucleotide database and the match with the highest similarity score was selected (http://blast.ncbi.nlm.nih.gov/Blast.cgi). Multiple nucleotide sequences representing all of the known species and subspecies of *Photorhabdus* and *Xenorhabdus* spp. were downloaded from the NCBI database (http://blast.ncbi.nlm.nih.gov/Blast.cgi), aligned with sequences from the study isolates, and trimmed to a 646 bp region using ClustalW^34^ in MEGA version 5.0^35^. Maximum likelihood trees were reconstructed using Nearest-Neighbor-Interchange (NNI) and Tamura-Nei model^36^ using MEGA version 5.05^35^. Bootstrap analysis was carried out with 1,000 datasets.

### Metabolite extraction

Bacterial cultures were grown in either SF900 media or Lysogeny broth (LB) for 72 hours at 30°C. A 1mL sample was taken from each culture and extracted with an equal volume of methanol, mixed briefly by vortexing and centrifuged for 30 minutes. The resulting supernatant was dried under a constant stream of nitrogen gas, to completion. Prior to measurement, samples were resuspended in 500 μL of methanol and centrifuged for 30 minutes.

### Ultra-performance liquid chromatography high-resolution mass spectrometry (UPLC-HRMS) measurements

UPLC-ESI-HRMS/MS analyses were performed using an UltiMate 3000 system linked to a Bruker Impact II qTof mass spectrometer. Runs were performed using a flow rate of 0.4 mL min^−1^ and gradient of MeCN/0.1% formic acid in H_2_O (5:95% to 95:5% over 15 mins). Data acquisition was performed as previously described^4^.

### Molecular Networking Analysis

The raw MS data of 114 environmental isolates, *E. coli* (all in LB and SF900), LB, SF900 and acetonitrile blanks were converted to the .mzXML format using DataAnalysis v4.3 (Bruker). Molecular networks were created using the online workflow at Global Natural Product Molecular Networking Social (GNPS)^15^.

The data was then clustered with MS-Cluster with a parent mass tolerance of .05 Da and a MS/MS fragment ion tolerance of .01 Da to create consensus spectra. Further, consensus spectra that contained less than 2 spectra were discarded. A network was then created where edges were filtered to have a cosine score above 0.7 and more than 6 matched peaks. Further edges between two nodes were kept in the network if and only if each of the nodes appeared in each other’s respective top 7 most similar nodes. The spectra in the network were then searched against GNPS’ spectral libraries. All matches kept between network spectra and library spectra were required to have a score above .7 and at least 6 matched peaks. Analog search was enabled against the library with a maximum mass shift of 100.0 Da. The self-loop networks were imported into Cytoscape (v3.4.0) for visualization.

### Feature identification

Mass spectrometry files were imported into DataAnalysis (v4.3) and converted from the Bruker .m format to the open mzXML format for processing with MZMine2^12^. After import, mass detection was performed with the mass detector set to centroid, noise level to 1000, at MS level 1 and with a retention time of 0-16.05 minutes. Chromatograms were then built with the retention time between 0-16.05 minutes, MS level 1, a minimum time span of 0.02, a minimum height of 1000 and an m/z tolerance of 0.005 m/z or 5.0 ppm. Peak deconvolution was performed with the noise amplitude algorithm, a minimum peak height of 1000, peak duration in the range 0-0.8 minutes and an amplitude of noise set to 5000.

The peak aligner was then set with an m/z tolerance of 0.005 m/z or 5.0 ppm, the weight of m/z at 20, retention time tolerance at 3% relative, weight for retention time of 10, with peaks requiring the same charge state, and ‘compare isotope pattern’ set to yes with the setting for isotope m/z tolerance 0.005 m/z or 5.0 ppm, a minimum absolute intensity of 1000, and a minimum score of 65%. Gap filling was then used using the ‘same RT and m/z range gap filler’ with m/z tolerance set to 0.005 m/z or 5.0 ppm. The aligned, filled mass list was then exported as a .csv file.

### Machine learning data pre-processing

In order to determine the importance of compounds, we decided to employ a machine learning model. In conjunction with a recently-developed feature attribution method, this serves the two-fold purpose of achieving a very high performance in discriminating between the two genera, yielding a model that can be subsequently used to classify new data, while at the same time allowing for a direct visualization of the features that have the largest impact on the model’s predictions for each of the samples.

The intensity and AUC data obtained from the MZmine2 peak picking algorithm were used. As a first step, we generated an additional dataset by setting to zero all AUC entries for which the corresponding peak intensity was zero. Samples were further processed by removing all columns corresponding to metabolites that were absent in all of the samples after deletion of *E. coli*, media only and acetonitrile blanks, since they would not contribute to the classification. In addition to this, we removed all columns with less than ca. 10% of non-zero values. The data were further cleaned up by clustering the metabolite columns according to their correlation across samples and discarding all but one of the members of any one cluster; the correlation thresholds used were 0.9, 0.95 and 0.99. Numerical metadata was scaled between 0 and 1 for pH, temperature and moisture, while the elevation, spanning three orders of magnitude, was converted to logarithmic scale. Location data, in turn, was kept to the level of province and one-hot-encoded; soil type and medium data was also one-hot-encoded. The smallest resulting dataset consisted of 20,650 and 21,634 metabolite columns, out of a total of 44,836, for the intensity and zeroed AUC data, respectively, plus 20 metadata columns: 2 media conditions, 4 soil types, 10 provinces, pH, temperature, moisture and elevation.

### Generating a model

The pruned datasets from the previous section were used to train a gradient boosting decision tree (GBDT) model. Here, we used the Python implementation of LightGBM^37^ to train a classifier on the pruned intensity and AUC datasets. We used 250 iterations, with 50 iterations as the threshold for early stopping, defined as the number of steps the model can take without improvements on the evaluation metric. The latter is calculated from the predictions of the model for a pre-defined validation set. To this end, we performed 100 rounds of 5-fold cross-validation on the datasets, and report the resulting mean and standard deviation of the mean accuracy and ROC-AUC (receiver operating characteristic curve - area under the curve) across folds.

### Determining feature importance

In order to interpret the predictions from the GBDT model and determine the most important features driving its output, we computed the SHAP values for each feature and averaged them over all the training rounds. The values are individualized per sample and correspond to the change in log-odds of the sample being classified as corresponding to one or the other genus – in this case, a positive value indicates a larger probability of being *Xenorhabdus* – relative to the mean prediction upon addition of a given feature, effectively measuring the impact that every feature value has on every sample. This was carried out using the tree ensemble implementation of the shap Python package^27^.

All code used for this paper is available at https://github.com/systemsmedicine/geographical-chemotypes as Jupyter notebooks, providing a step-by-step walkthrough.

### Compound isolation and purification

For the isolation and purification of (4*R*,8*R*,12*R*,16*R*)-4,8,12,16-tetramethyl-1,5,9,13-tetraoxacyclohexadecane-2,6,10,14-tetrone, the XAD-16 resin from a 4 L M63 medium culture of *X. szentirmaii_*P1 (phenazine gene cluster knockout) mutant^38^ were harvested after 72 h of incubation at 30°C with shaking at 120 rpm, washed with water and extracted with methanol (3 × 1 L) to yield the crude extract (1.1 g) after evaporation. The extract was dissolved in methanol and was subjected to preparative HPLC-MS with C-18 column (21.2 mm × 250 mm, 7.0 *µ*m, Agilent) using an acetonitrile/water gradient (0.1% formic acid) in 30 min, 5-95% to afford a sub-fraction mainly containing 8.3 mg. The sub-fraction was further purified by semipreparative HPLC with C-18 column (9.4 mm × 250 mm, 5.0*µ*m, Agilent) using an acetonitrile/water gradient (0.1% formic acid) 0-30 min, 30-45% to afford (4*R*,8*R*,12*R*,16*R*)-4,8,12,16-tetramethyl-1,5,9,13-tetraoxacyclohexadecane-2,6,10,14-tetrone (2.1 mg). ^1^H and ^13^C NMR, ^1^H-^13^C Heteronuclear Single Quantum Coherence (HSQC), ^1^H-^13^C Heteronuclear Multiple Bond Correlation (HMBC), and ^1^H-^1^H Correlation Spectroscopy (COSY) were measured. Chemical shifts (δ) were reported in parts per million (ppm) and referenced to the solvent signals. Data are reported as follows: chemical shift, multiplicity (d = doublet, dd = doublet of doublet, and m = multiplet), and coupling constants in Hertz (Hz).

## Acknowledgements

The authors would like to thank Dr. Lothar Fink from Goethe University for conducting the X-ray crystallography structure determination. Financial support was provided by Naresuan University (Grant Number R2560B073). CPR and EAHV were supported by the Alfons und Gertrud Kassel-Stiftung. YMS is the recipient of a Humboldt Postdoctoral Fellowship. Work in the Bode lab was supported by the LOEWE-TBG initiative.

