## Supplemental material for "Focused natural product elucidation by prioritizing high-throughput metabolomic studies with machine learning"

**Supplementary Material**

**Supplementary Figures**

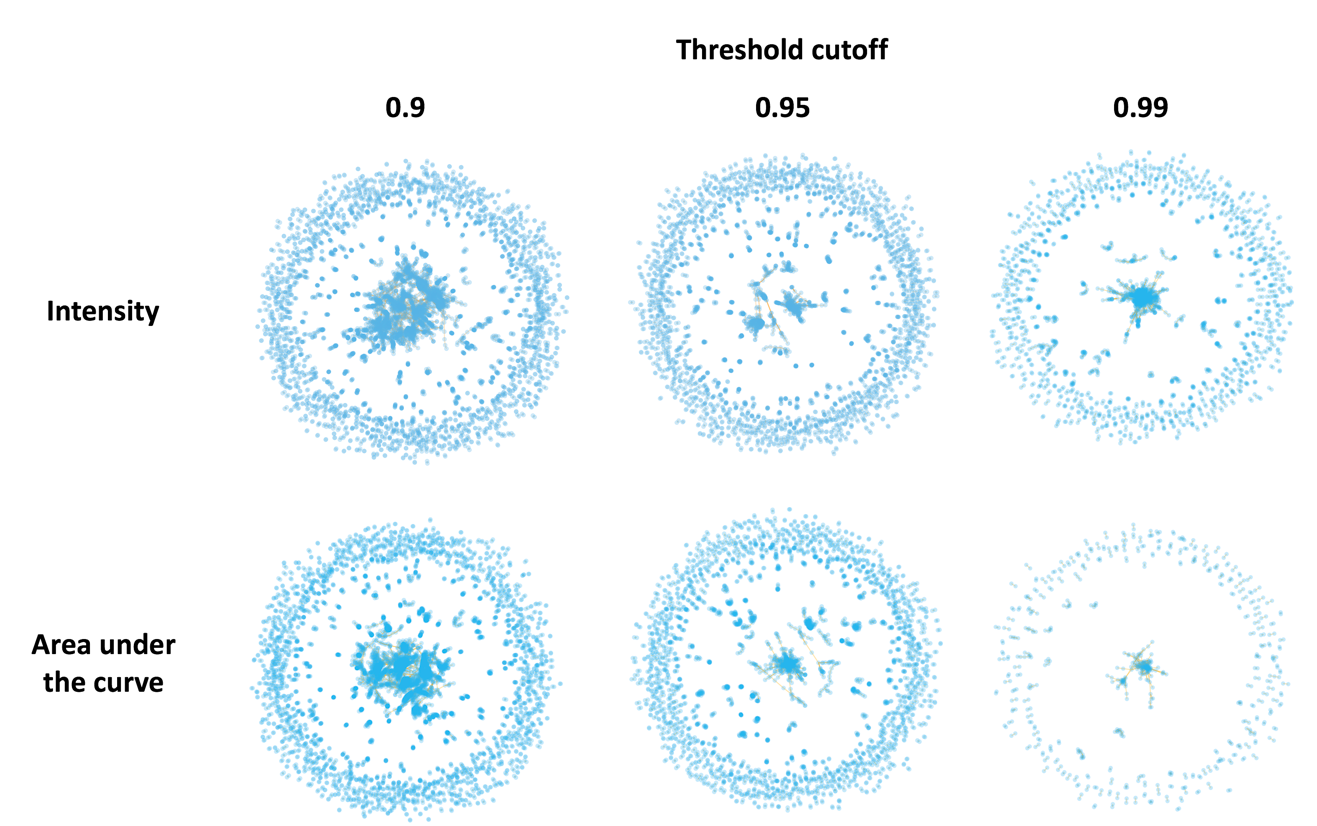

**Supplementary Figure S1.** Clustering of features of either intensity or AUC was performed using three different correlation cutoffs; 0.9, 0.95 and 0.99. These clusters can be explored interactively at [cparrarojas.github.io/blog/2019/02/geographical-chemotypes](https://cparrarojas.github.io/blog/2019/02/geographical-chemotypes/).

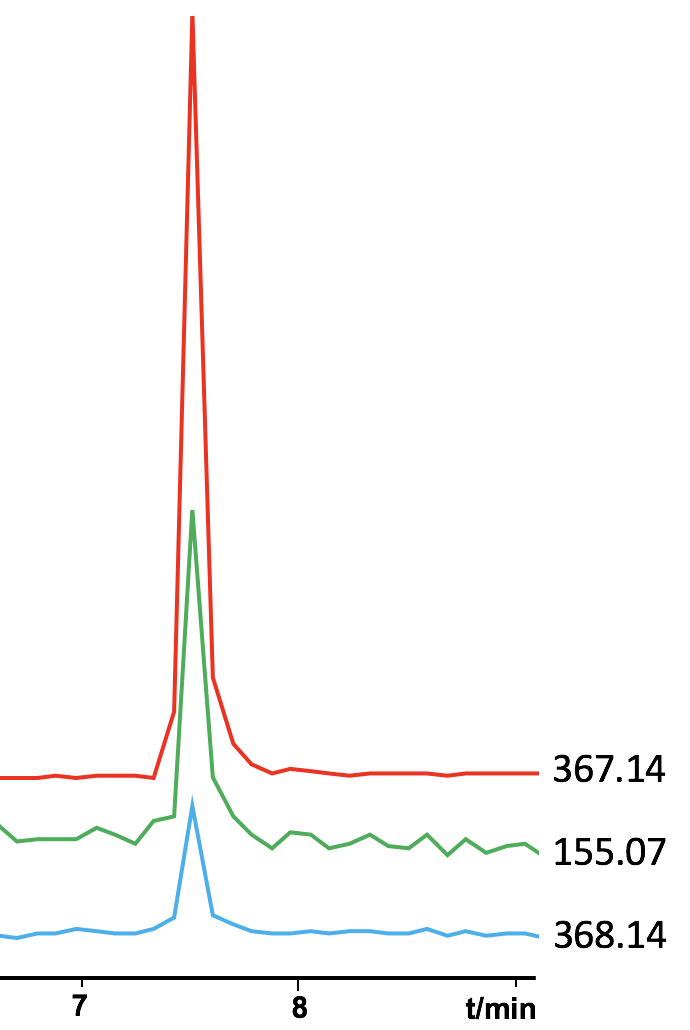

**Supplementary Figure S2.** Representative extracted ion chromatograms of the top 3 ranking features as determined by the machine learning model.

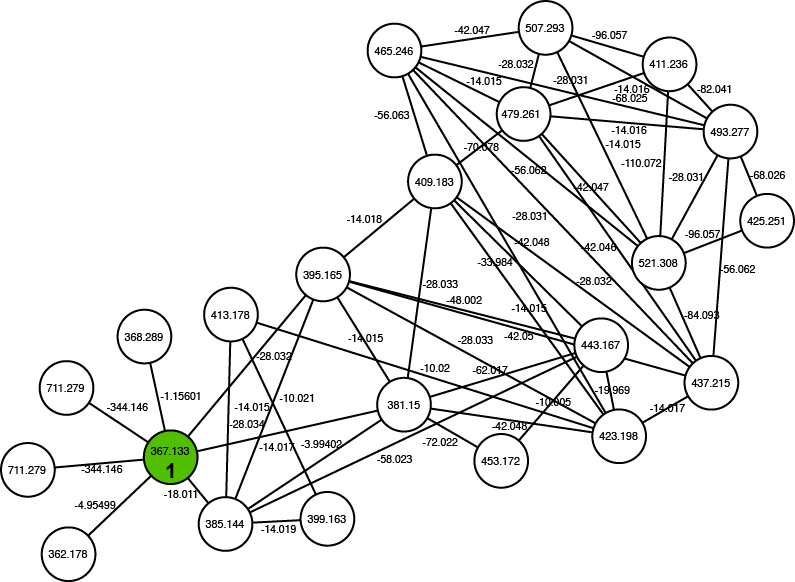

**Supplementary Figure S3.** Network containing **1** showing parent masses inside nodes and edges representing mass differences between metabolites.

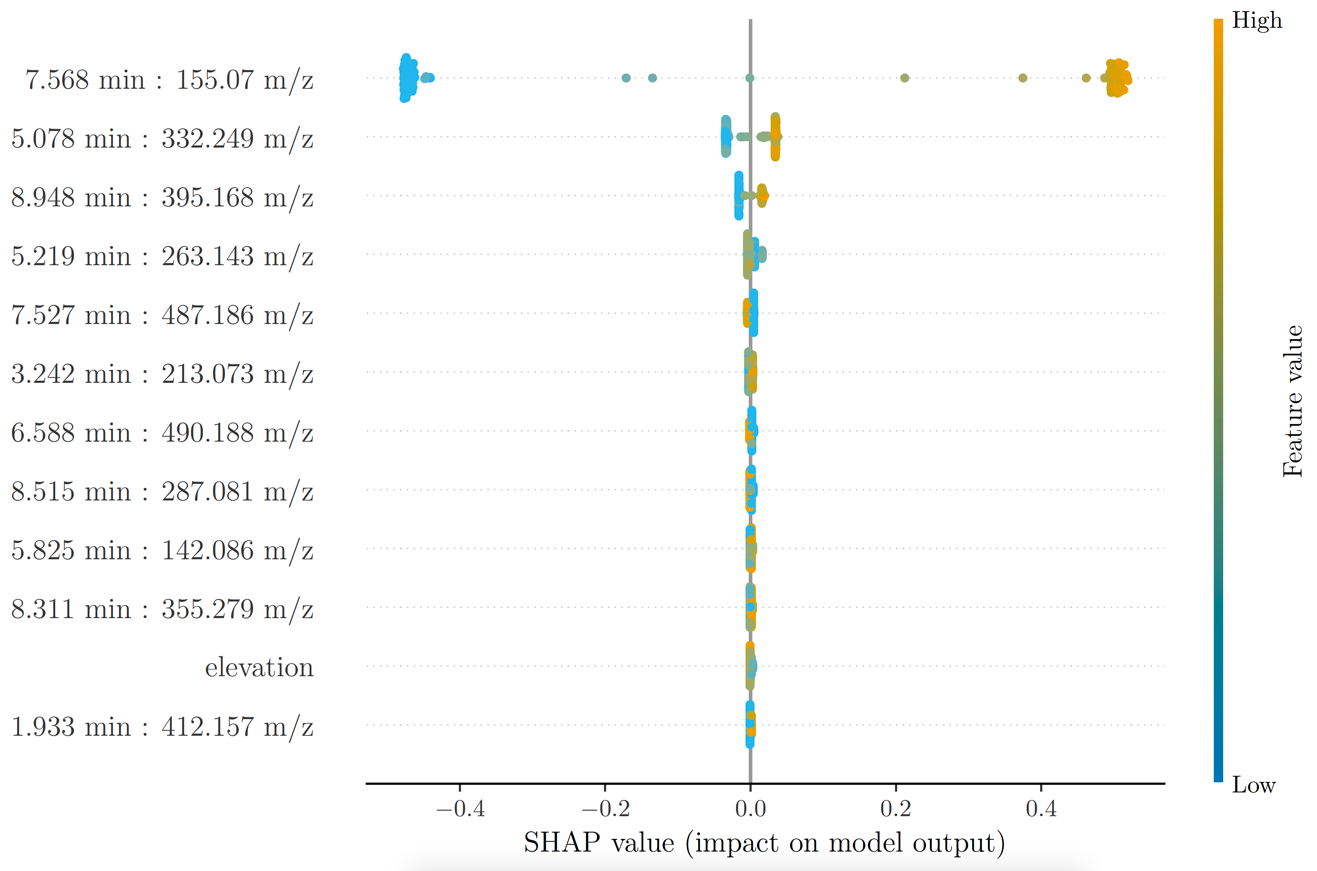

**Supplementary Figure S4.** SHAP values were generated on models following the clustering of intensity data based upon a correlation cut-off of 0.9.

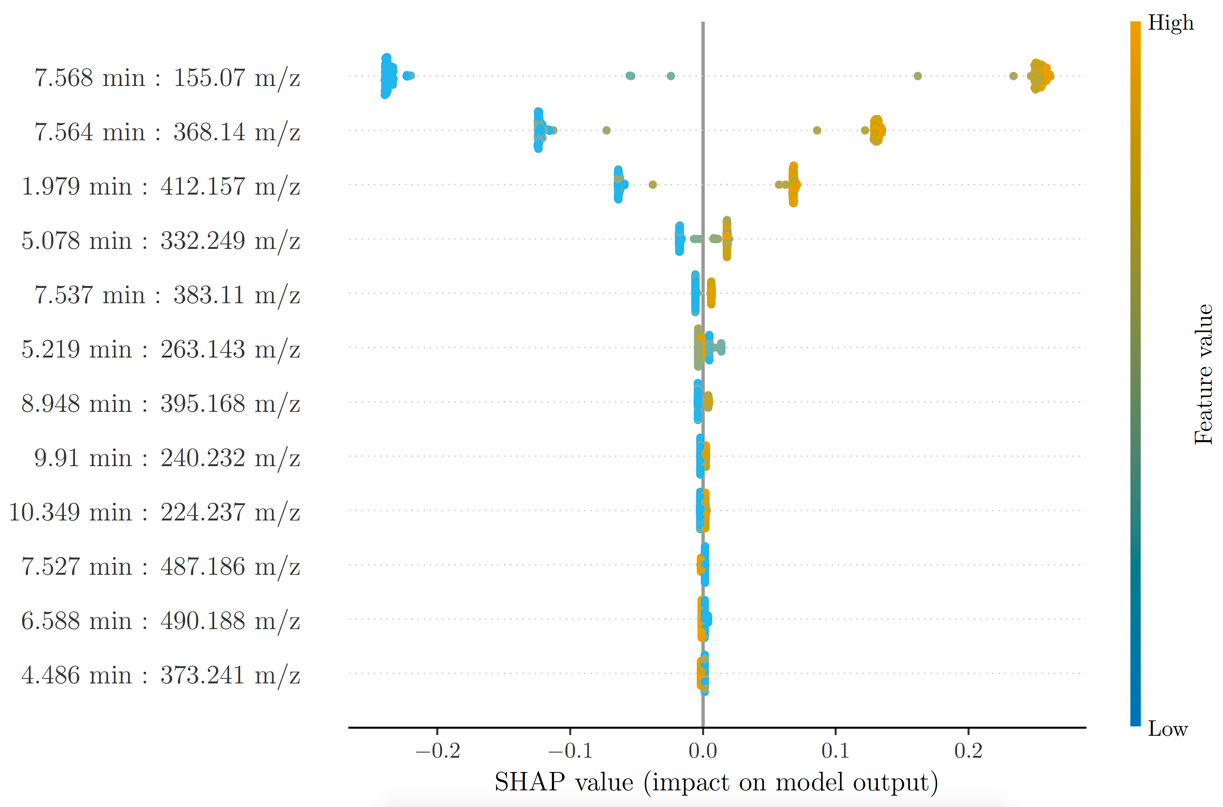

**Supplementary Figure S5.** SHAP values were generated on models following the clustering of intensity data based upon a correlation cut-off of 0.95.

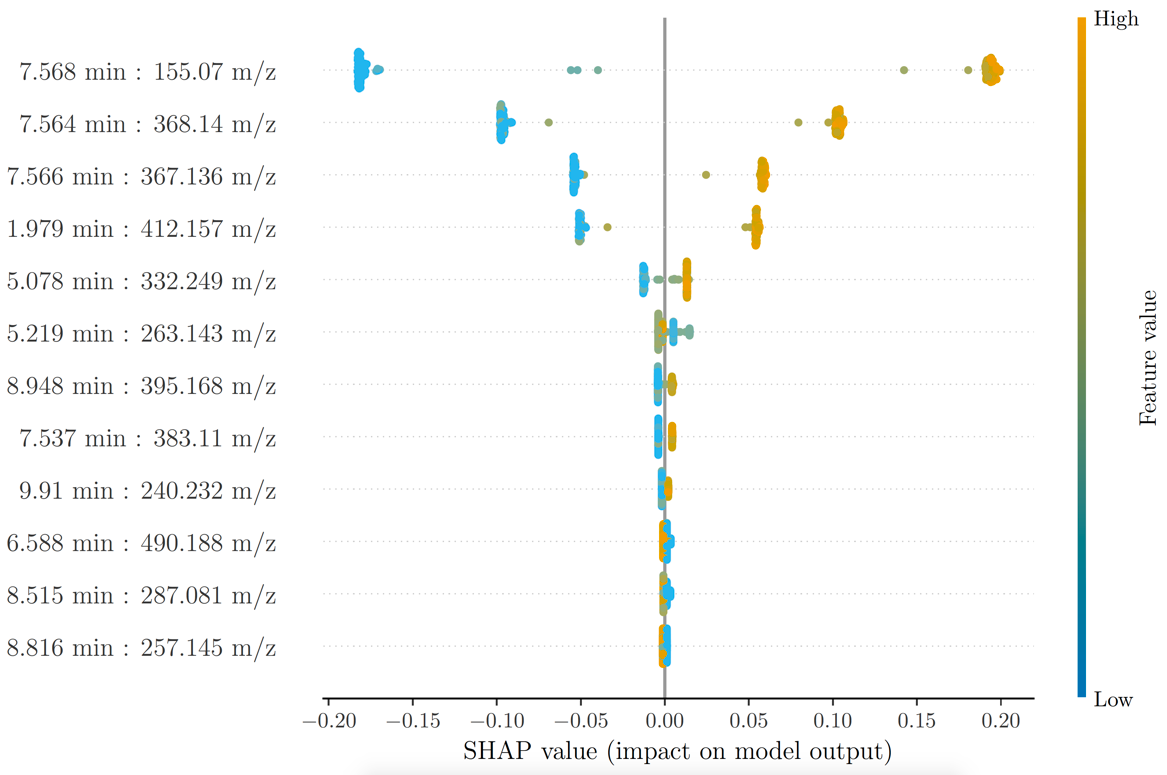

**Supplementary Figure S6.** SHAP values were generated on models following the clustering of intensity data based upon a correlation cut-off of 0.99.

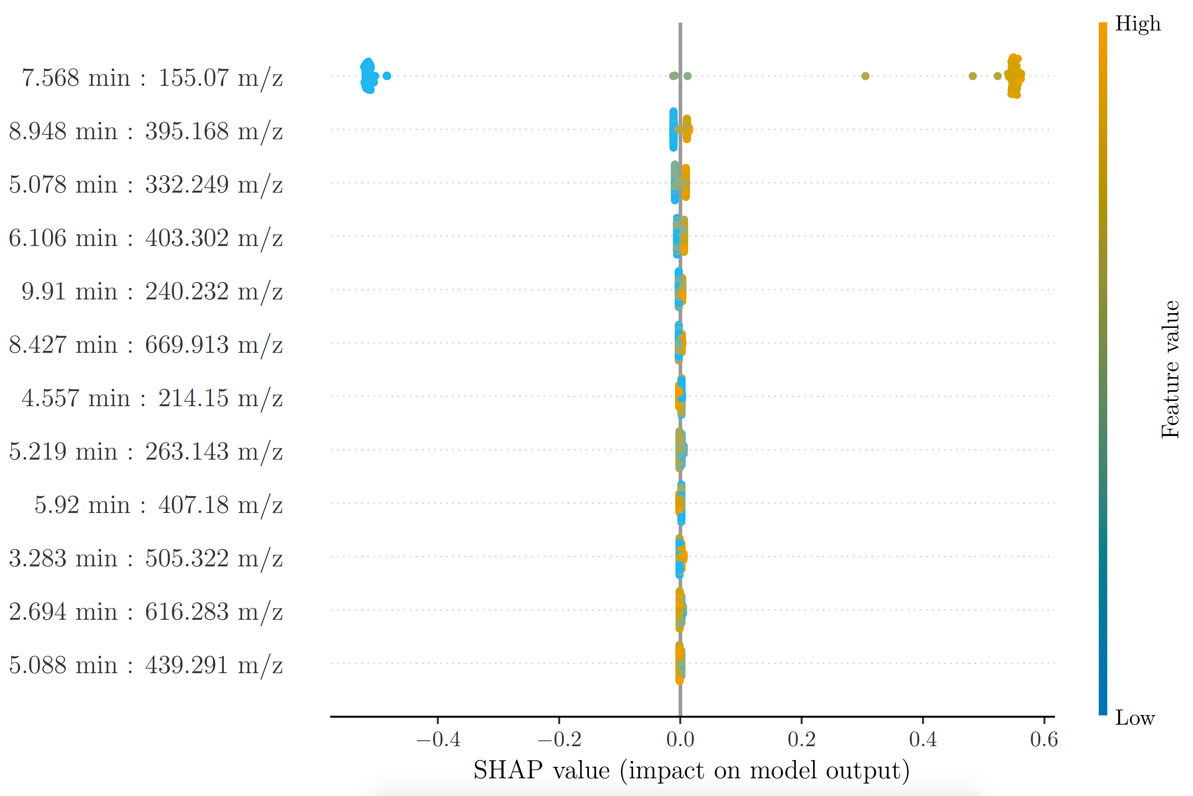

**Supplementary Figure S7.** SHAP values were generated on models following the clustering of AUC data based upon a correlation cut-off of 0.9.

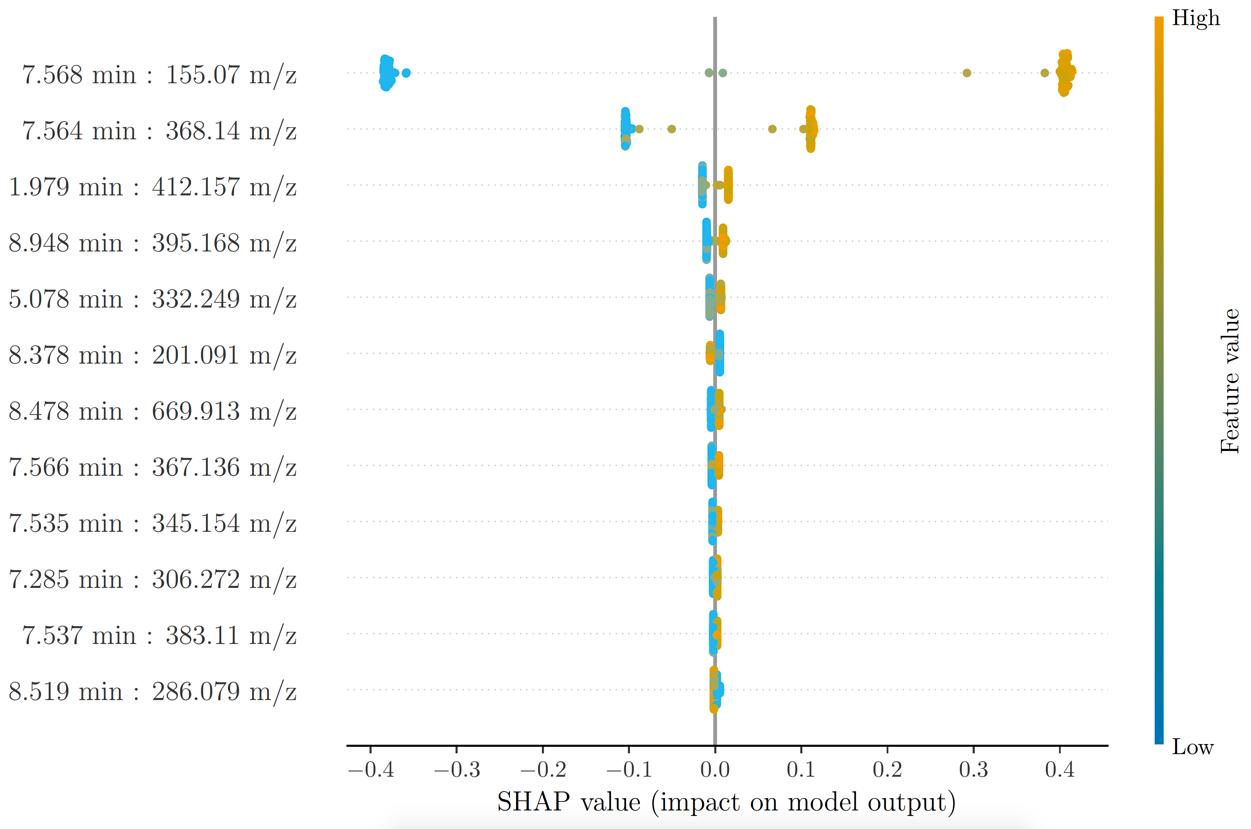

**Supplementary Figure S8.** SHAP values were generated on models following the clustering of AUC data based upon a correlation cut-off of 0.95.

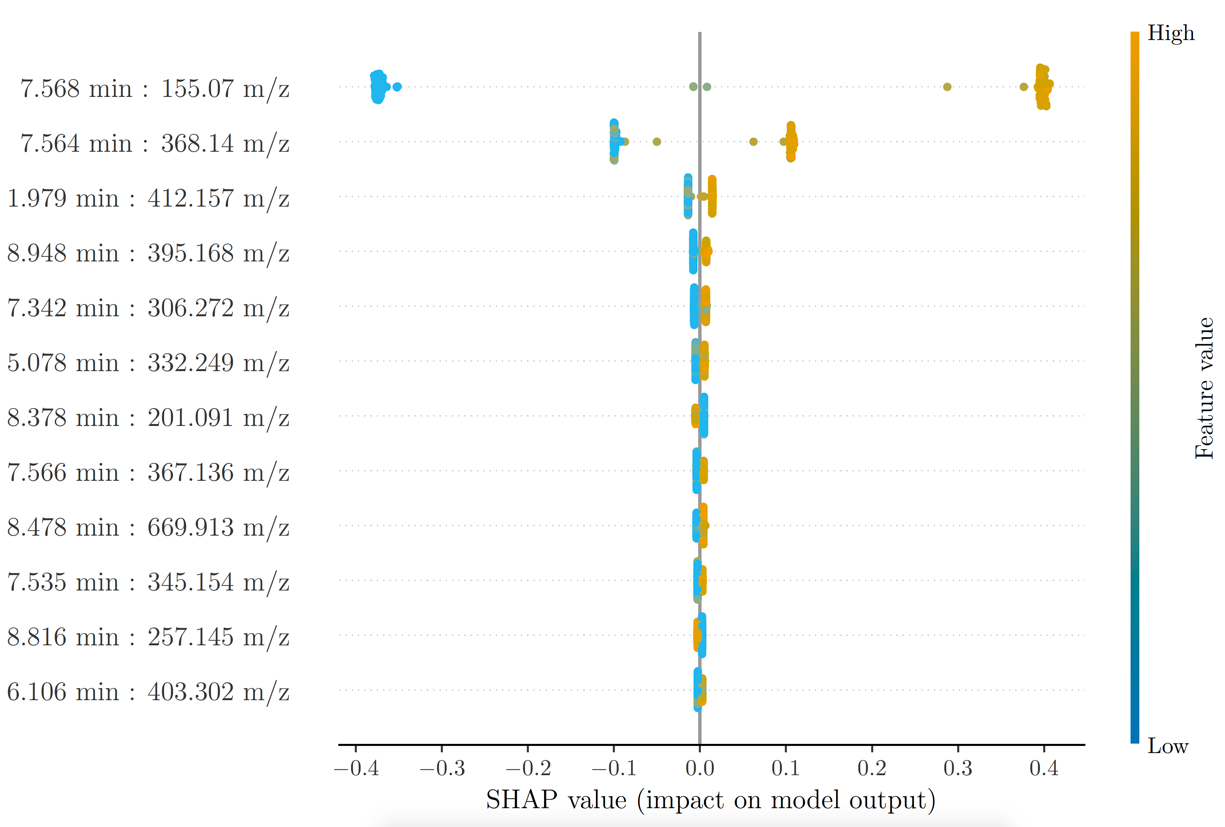

**Supplementary Figure S9.** SHAP values were generated on models following the clustering of AUC data based upon a correlation cut-off of 0.99.

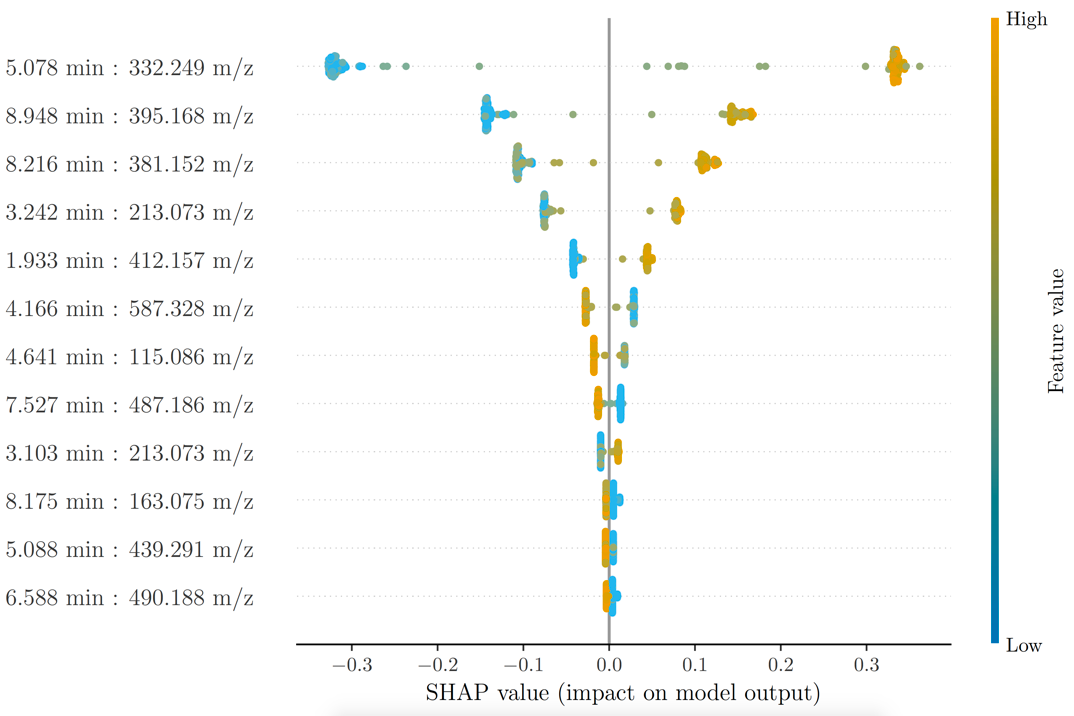

**Supplementary Figure S10.** All features associated with the top-ranking cluster were removed and the model was recalculated. The top 10 highest ranking features are shown based on the output of SHAP. Performance metrics can be seen in Supplementary Table 4.

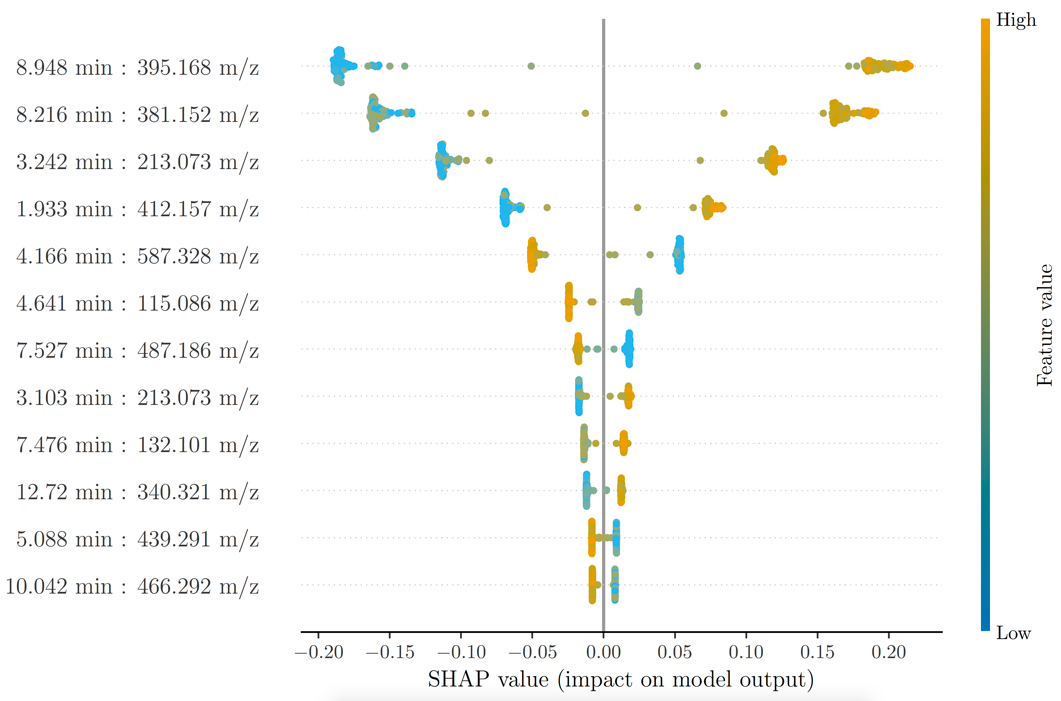

**Supplementary Figure S11.** All features associated with the top two ranking clusters were removed and the model was recalculated. The top 10 highest ranking features are shown based on the output of SHAP. Performance metrics can be seen in Supplementary Table 4.

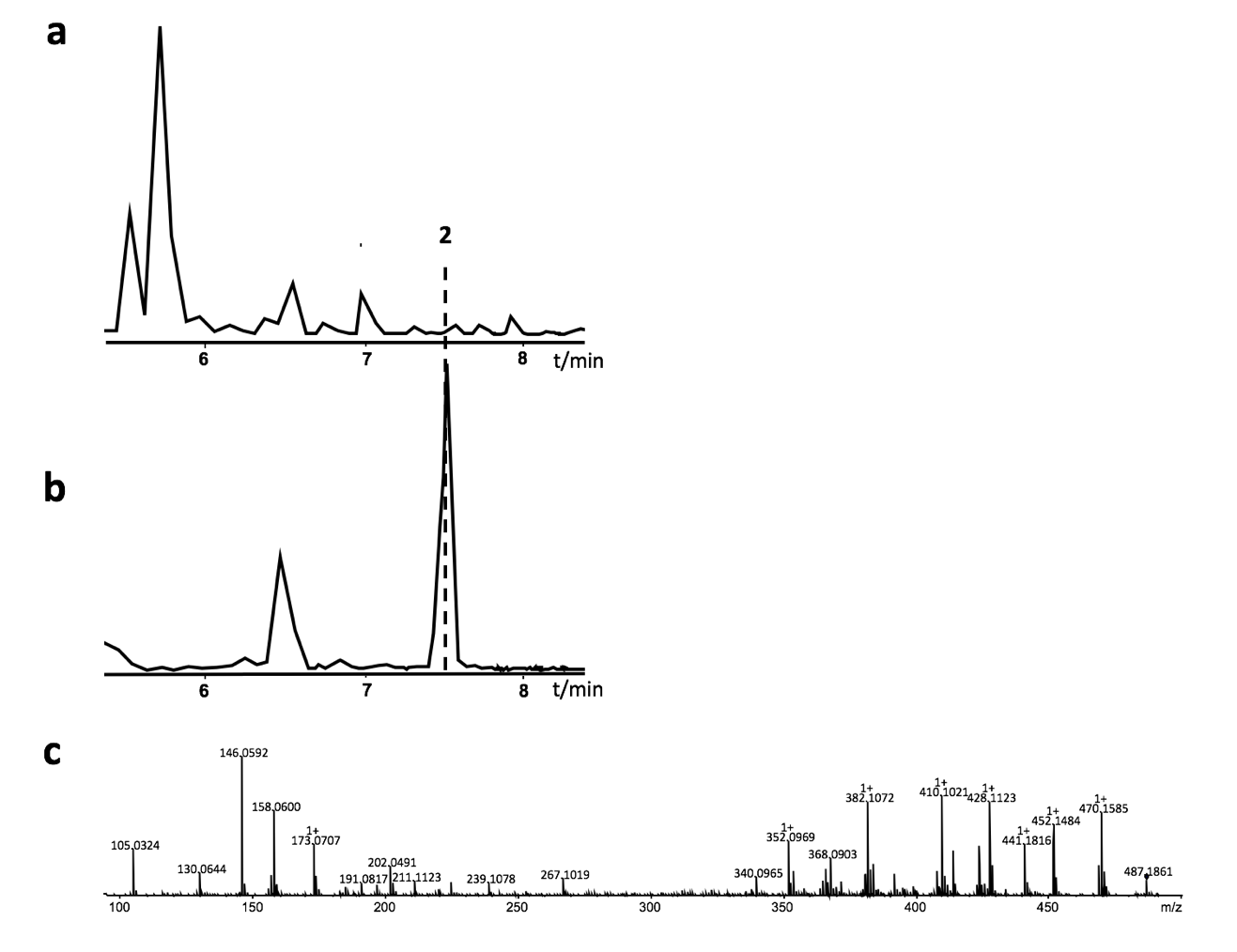

**Supplementary Figure S12a.** Base peak chromatogram of a representative *Photorhabdus* strain (number 448), **(b)** with an extracted ion chromatogram of the signal with the highest negative correlation to **1** and *m/z* of 487.186. **(c)** The fragmentation pattern of this compound, **2**, is also shown.

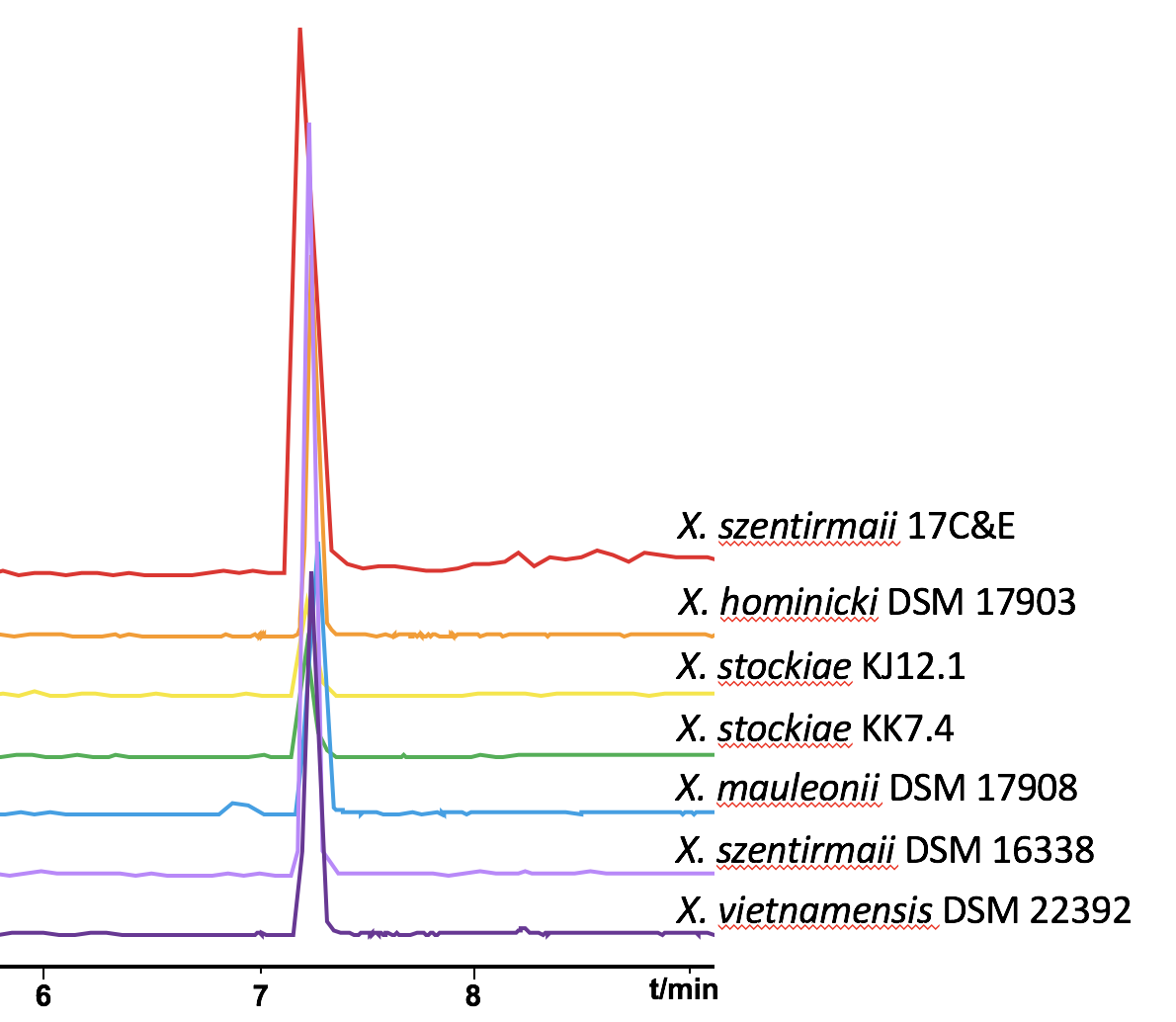

**Supplementary Figure S13.** Extracted ion chromatograms of other *Xenorhabdus* species containing the (cyclo)tetrahydroxybutyrate (**1**).

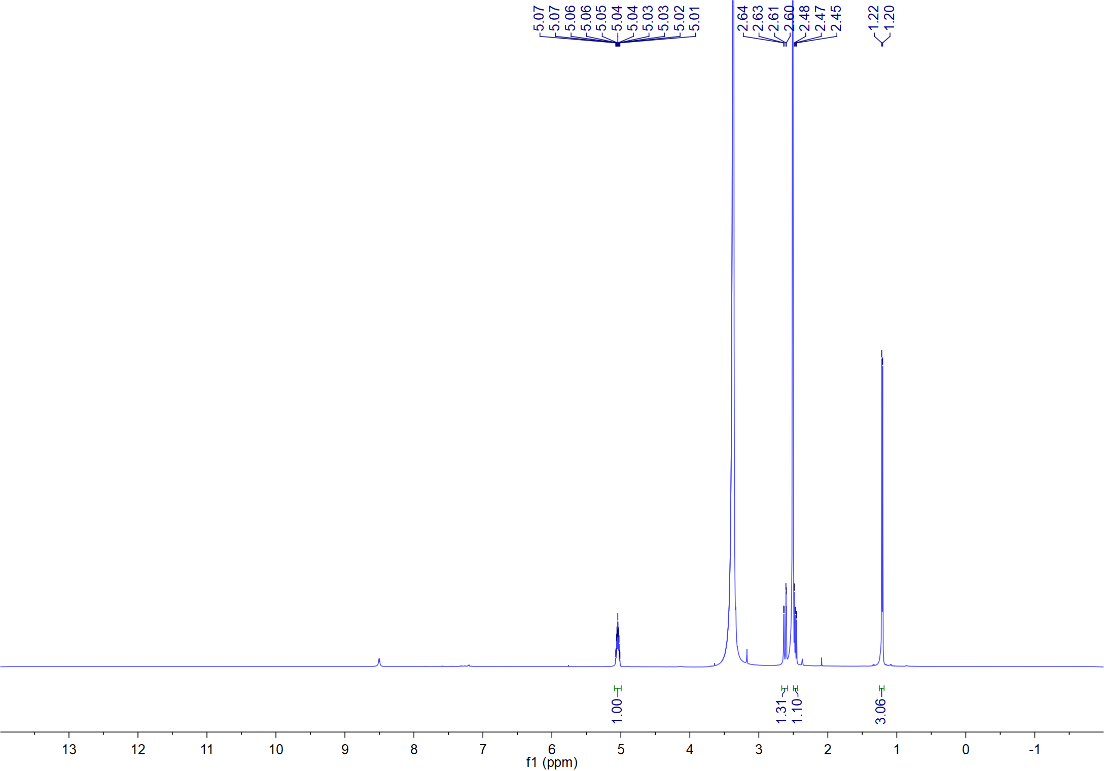

**Supplementary Figure S14.** ^1^H NMR spectrum of **1** in DMSO-*d*_6_.

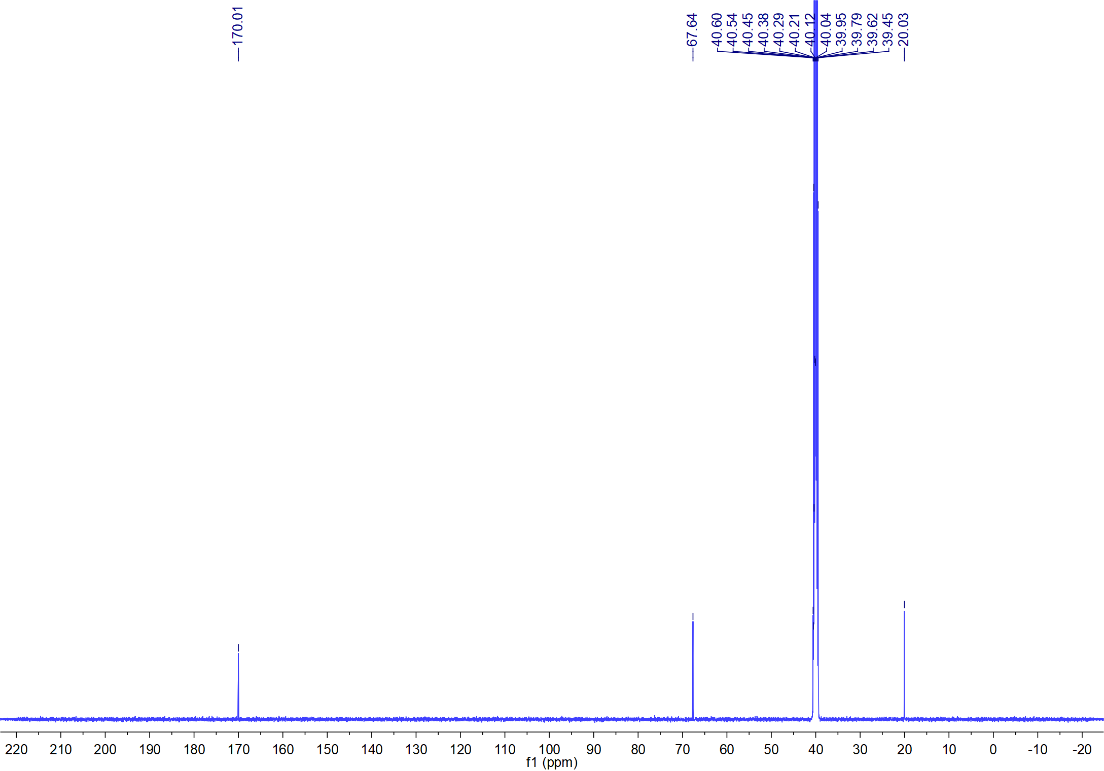

**Supplementary Figure S15.** ^13^C NMR spectrum of **1** in DMSO-*d*_6_.

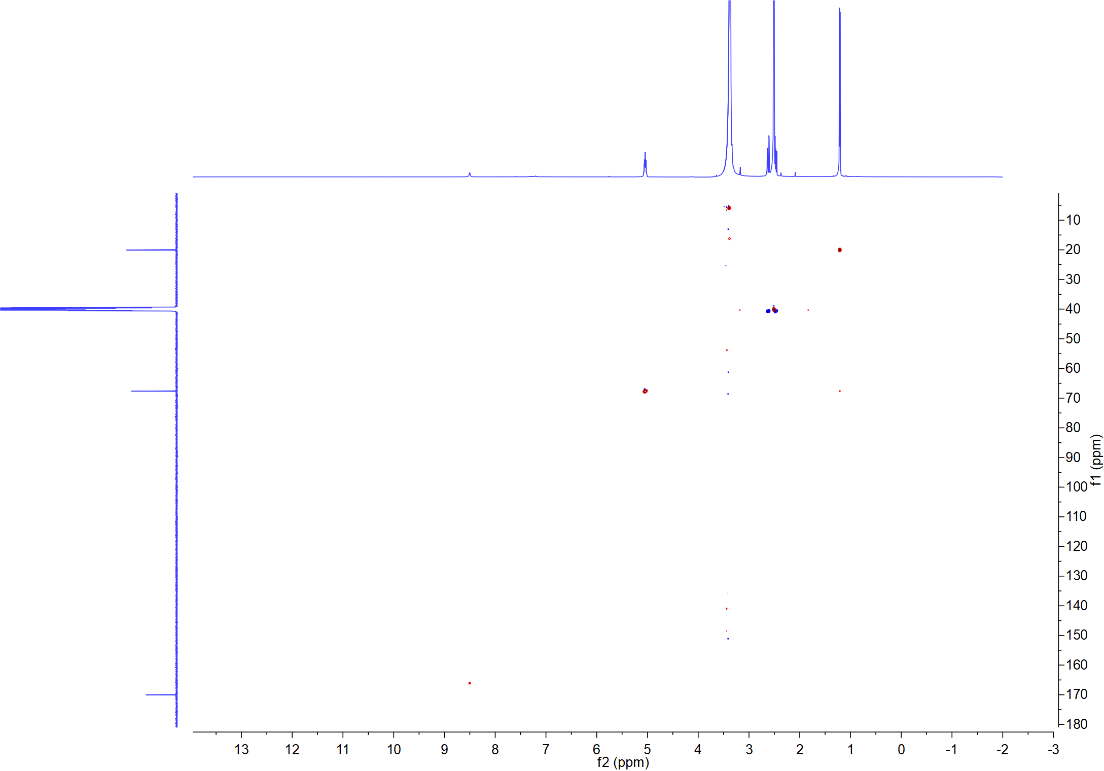

**Supplementary Figure S16.** HSQC spectrum of **1** in DMSO-*d*_6_.

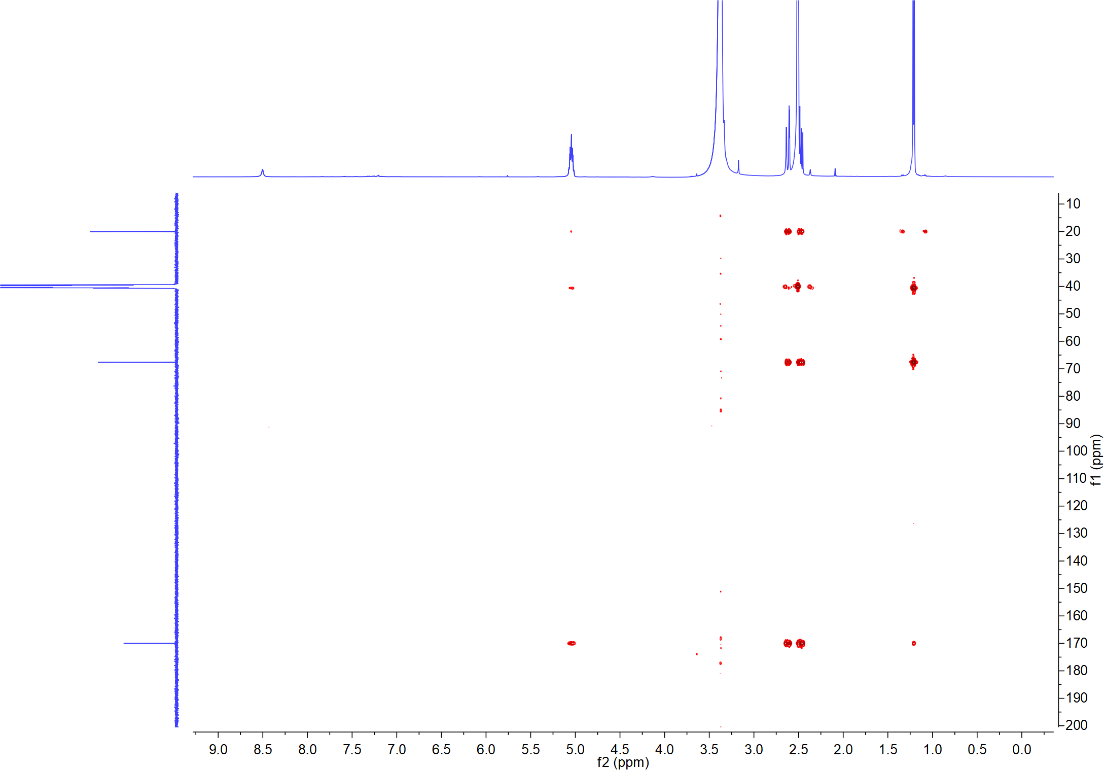

**Supplementary Figure S17**. HMBC spectrum of **1** in DMSO-*d*_6_.

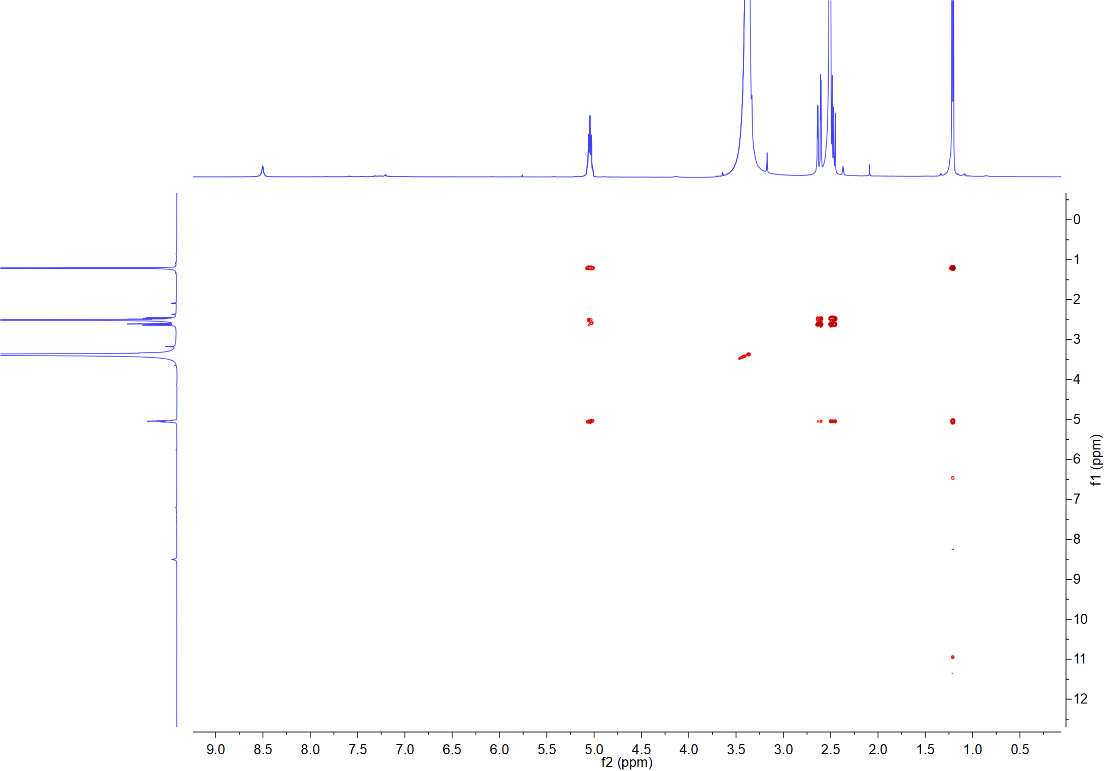

**Supplementary Figure S18.** ^1^H-^1^H COSY spectrum of **1** in DMSO-*d*_6_.

**Supplementary Tables**

**Supplementary Table S2.** Performance of gradient boosting decision tree model on full data set compared to that of the pruned and clustered data. Clustered data is shown with different cutoff thresholds. All models were calculated using both intensity data and area under the curve (AUC). The degree of clustering can be seen in Supplementary Figure S1.

|  | **Full data set** | | **Highly-correlated & clustered data** | | | | | |
| --- | --- | --- | --- | --- | --- | --- | --- | --- |
|  |  |  | **0.9** | | **0.95** | | **0.99** | |
|  | **Intensity** | **AUC** | **Intensity** | **AUC** | **Intensity** | **AUC** | **Intensity** | **AUC** |
| **Accuracy** | 97.6% | 97.4% | 97.5% | 97.3% | 97.5% | 97.4% | 97.6% | 97.4% |
| **Standard deviation** | 0.67% | 0.80% | 0.67% | 0.91% | 0.67% | 0.77% | 0.67% | 0.80% |

### **Supplementary Table S3.** ^1^H (500 MHz) and ^13^C (125 MHz) NMR data assignments for **1** in DMSO-*d*_6_ (for NMR spectra see Figure S14-S18).

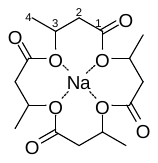

| No. | *δ*_H_ (mult., *J*) | *δ*_C_ |
| --- | --- | --- |
| 1 | - | 170.0, C |
| 2 | 2.62 (dd, 16.1, 2.7) | 40.60, CH_2_ |
|  | 2.47 (overpal) |  |
| 3 | 5.04 (m) | 67.6, CH |
| 4 | 1.21 (d, 6.4) | 20.0, CH_3_ |

**Supplementary Table S4.** Performance of model when using previously identified highly-ranked signals as a single predictor.

|  | **Intensity** | | | **AUC** | | |
| --- | --- | --- | --- | --- | --- | --- |
| ***m/z*** | **155.07** | **368.14** | **367.14** | **155.07** | **368.14** | **367.14** |
| **Accuracy** | 97.5% | 97.9% | 97.6% | 97.4% | 97.5% | 96.4% |
| **Standard deviation** | 0.48% | 0.50% | 0.43% | 0.36% | 0.60% | 0.62% |

**Supplementary Table S5.** Model performance of single features as sole predictors on unseen data. A total of 15 *Xenorhabdus* (X) and 14 *Photorhabdus* (P) were grown in triplicate and their metabolites extracted as described in the Methods. Listed are percentages representing how often the correct genus was called. Probabilities for calling each sample can be seen in Supplementary Table S6.

|  | **Intensity** | | | | | | **Area under the curve** | | | | | |
| --- | --- | --- | --- | --- | --- | --- | --- | --- | --- | --- | --- | --- |
| ***m/z*** | **155.07** | | **368.14** | | **367.14** | | **155.07** | | **368.14** | | **367.14** | |
|  | LB | SF900 | LB | SF900 | LB | SF900 | LB | SF900 | LB | SF900 | LB | SF900 |
| ***P*** | 100 | 100 | 100 | 100 | 100 | 100 | 92.9 | 95.2 | 100 | 100 | 100 | 97.6 |
| ***X*** | 84.4 | 91.1 | 88.9 | 91.1 | 88.9 | 91.1 | 93.3 | 95.2 | 93.3 | 91.1 | 93.3 | 93.3 |
| **Overall** | 92.0 | 95.4 | 94.3 | 95.4 | 94.3 | 95.4 | 93.1 | 95.4 | 96.6 | 95.4 | 96.6 | 95.4 |
